# FOntCell: Fusion of Ontologies of Cells

**DOI:** 10.1101/850131

**Authors:** Javier Cabau-Laporta, Alex M. Ascensión, Mikel Arrospide-Elgarresta, Daniela Gerovska, Marcos J. Araúzo-Bravo

## Abstract

High-throughput cell-data technologies such as single-cell RNA-Seq create a demand for algorithms for automatic cell classification and characterization. There exist several classification ontologies of cells with complementary information. However, one needs to merge them in order to combine synergistically their information. The main difficulty in merging is to match the ontologies since they use different naming conventions. To overcome this obstacle we developed an algorithm that merges ontologies by integrating the name-matching search between class label names with the structure mapping between the ontology elements. To implement our algorithms, we developed FOntCell, a software module in Python for efficient automatic parallel-computed fusion of ontologies in the same or similar knowledge domains. It processes the ontology attributes to extract relations and class synonyms. FOntCell integrates the semantic, name with synonyms, mapping with a structure mapping based on graph convolution. Since the structure mapping assessment is time consuming process, we designed two methods to perform the graph convolution: vectorial structure matching and constraint-based structure matching. To perform the vectorial structure matching we designed a general method to calculate the similarities between vectors of different lengths for different metrics. Additionally, we adapted the slower Blondel method to work for structure matching. These functionalities of FOntCell allow the unification of dispersed knowledge in one domain into a unique ontology. FOntCell produces the results of the merged ontology in OBO format that can be iteratively reused by FOntCell to adapt continuously the ontologies with the new data, such of the Human Cell Atlas, endlessly produced by data-driven classification methods. To navigate easily across the fused ontologies, it generates HTML files with tabulated and graphic summaries, and an interactive circular Directed Acyclic Graphs of the merged results. We used FOntCell to fuse CELDA, LifeMap and LungMAP Human Anatomy cell ontologies to produce comprehensive cell ontology.

**Author Summary:** There is a strong belief in the research community that there exist more cell types than the described in the literature, therefore new technologies were developed to produce a high volume of data to discover new cells. One issue that arises once the cells are discovered is how to classify them. One way to perform such classification is to use already existing cell classifications from different ontology sources but it is difficult to merge them. An ontology has semantic information providing the meaning of each term and structural information providing the relationship between terms as a graph. We developed a new Python module, FOntCell that merges efficiently cell ontologies and integrates semantic and structure information with our own graph convolution technique. Since the structure mapping assessment is time-consuming process we designed two methods to optimize the graph convolution: vectorial and constraint-based structure matching. To perform the vectorial structure matching we designed a method that calculates the similarities between vectors describing the graphs of different sizes. The functionalities of FOntCell allow the unification of dispersed knowledge into a unique ontology, to adapt continuously from new data, and to navigate across the fused ontologies by a graphic use interface.

## Introduction

Precision biomedicine technologies produce a growing quantities of detailed information in the form of high throughput data from finer-grained biomedical samples reaching single-cell (1) and subcellular (2) levels. This increasingly precise cellular data make obsolete the existing cell classification systems and creates the demand for automatic comprehensive data-driven cell classification methods. Among the structures to classify knowledge domain items are the ontologies; they can be defined in several ways depending on the context of use (3). In information science, an ontology is defined as a seven-tuple, O:= {L, C, R, F, G, T, A}, where L:=LC u LR is a lexicon of concepts LC and relations LR; C is a set of concepts; R is a set of binary relations on C; F is a function connecting symbol concepts to sets of concepts Sub(LC)→Sub(C); G is a function connecting symbol relations to sets of relations, Sub(LR)→Sub(R); T is a taxonomy for the partial ordering of C, T(C_i_, C_j_), and A is a set of axioms with elements C and R (3). A critical question needing an answer during the design of ontology is the level of detail covered by the ontology. Thus, different ontologies of the same knowledge domain use different conceptualizations to obtain the desired level of granularity. In the case of cell ontologies, there are several cell type classifications in various formats; the most frequently used being the format of Open Biomedical Ontologies (OBO). Single-cell analysis is broadening the discovery of new cell types, prompting the need for advanced methods to classify these new cells as branches of the existing cell ontologies. Traditionally, such cell classification relies on human data curation. However, the growing number of cell types boosted by high throughput data generation such as single cell RNA-Seq creates a necessity to develop automatic computational methods assisting cell ontology creation (4). New cell ontologies can be created by reusing the information dispersed in multiple cell ontologies and merging it.

There exist numerous semiautomatic tools for alignment and merging of ontologies (Table 1). Tools like ATOM (6) are omitted since they are algorithms for ontology merging that do not introduce alignment methods and require mapping as input. The majority of the methods require an initial input and some intermediate user inputs for performing correct alignment thus they are semi-automatic methods. Nowadays, new cell discoveries are abundant thanks to single-cell RNA-Seq technologies and international research initiatives such as the Human Cell Atlas (HCA). To decrease the human involvement to a minimum in the merging of new cell ontologies, we developed an algorithm and implemented it as a software package in Python, FOntCell, for automatic fusion of ontologies. We applied FOntCell to create a new, more comprehensive and fine-grained cell ontology by merging cell ontologies, thus expanding automatically the hierarchical information of the component ontologies.

**Table 1.**
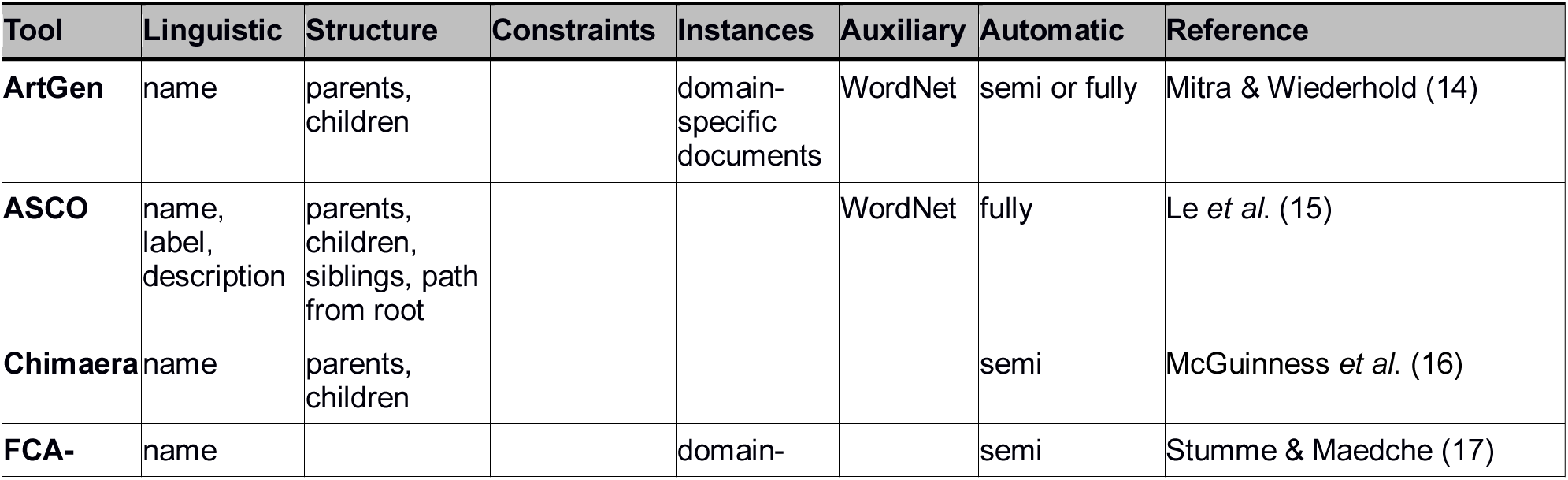
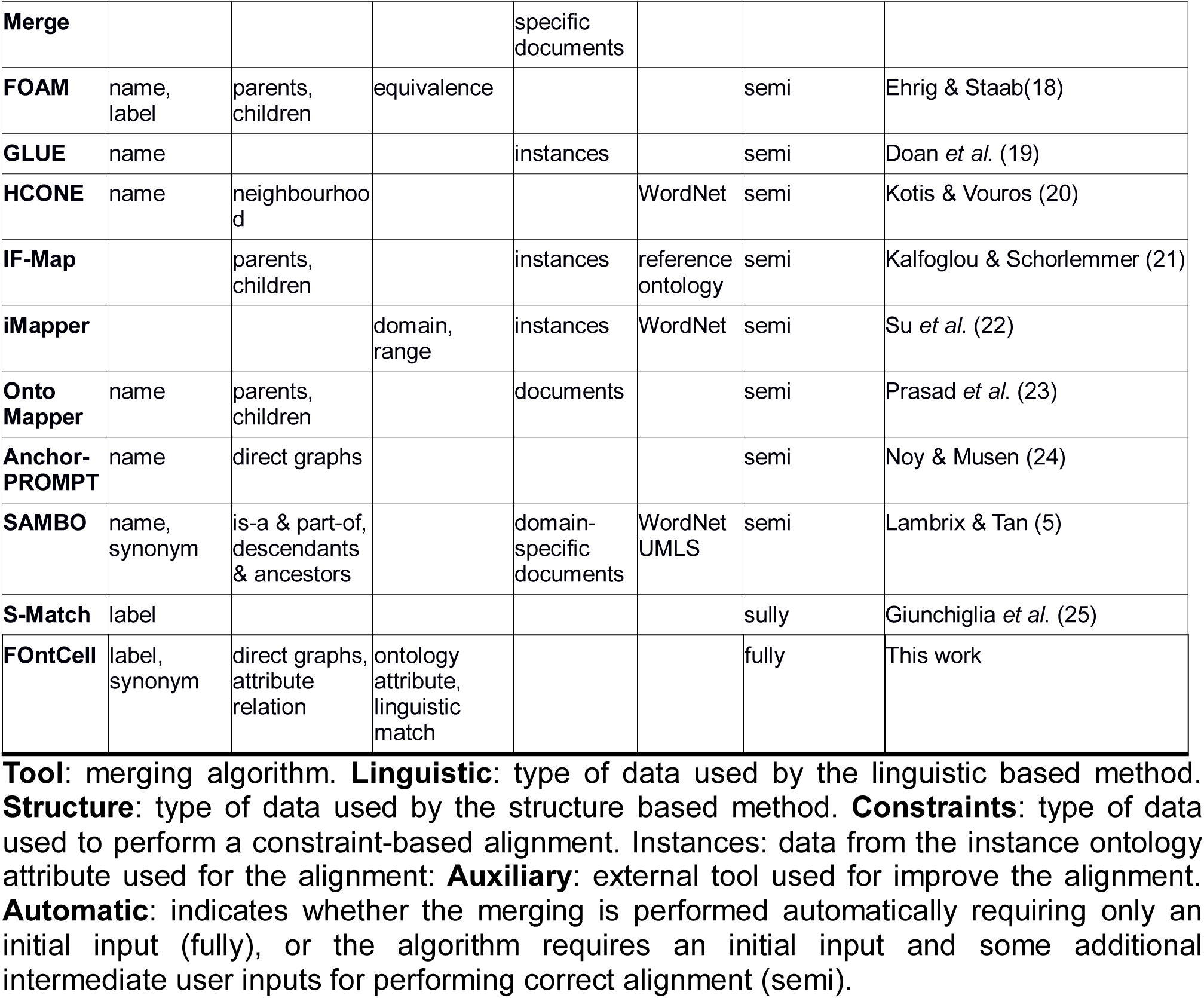
Tools for alignment and merging of ontologies and their features, adapted from Table 9 from Lambrix and Tan(5).

There are multiple ontologies with biomedical information (genomics, proteomics, and anatomy) (7). Two of the biggest cell type ontologies are CELDA (8) and LifeMap (9). CELDA integrates information about gene expression, localization, development and anatomy for *in vivo* and *in vitro* human and mouse cells, as well as cell development. With FOntCell, we aim to build a comprehensive and more specific ontology of the cellular development, giving rise to all cell types of the human body, which is highly needed for the characterization of the growing quantities of data produced by single-cell analysis in projects such as the HCA. Therefore, we focused on the ‘development’ annotation information of CELDA stored in the fields CL (Cell Ontology), CLO (Cell Line Ontology) and EFO (Experimental Factor Ontology). Another important repository for cell information is LifeMap (9); storing knowledge on *in vivo* cellular development, cell type and gene expression. LifeMap provides contrasted data and enough cell types to be fused synergistically with CELDA, both ontologies containing cell types missing in the other. An issue arising when fusing CELDA and LifeMap is their different labelling systems. The two ontologies use different labels to name same cell types, thus, a simple word matching cannot find equivalences. Therefore, it is necessary to align ontologies (10), setting the classes of one ontology equivalent to the classes of the other ontology. We developed an algorithm that can interpret two classes from two ontologies as equivalent, taking into account not only the class labelling but also the internal structure of the ontologies.

## Results

### The merging of CELDA with LifeMap produced a growth of CELDA by 67.42%

To find the optimal parameters of FOntCell for the merging of CELDA with LifeMap, we performed a bidimensional scanning of the parameters local name threshold *θ*_*LN*_ and window length *W* in the range [0.1, 0.8] and [1, 8], respectively, using steps of 0.1 for *θ*_*LN*_, and 1 for *W* for all structure mapping metrics: the three vectorial structure matching methods, namely Euclidean, Pearson, and cosine, the constraint-based structure matching, and Blondel structure matching (Fig.1A). We kept the name mapping threshold *θ*_*N*_ that provides good name matching results at 0.85. For all possible pairwise combinations of *θ*_*LN*_ and *W*, we observed a variation between the structure matching obtained by structure scores *T*^*AB*^(*i*,*j*) that pass the *θ*_*T*_ and *θ*_*LN*_ thresholds between ontology *A* = CELDA and *B* = LifeMap. The constraint-based method does not involve local name matching assessment, therefore *θ*_*LN*_ was not used.

**Figure 1.**
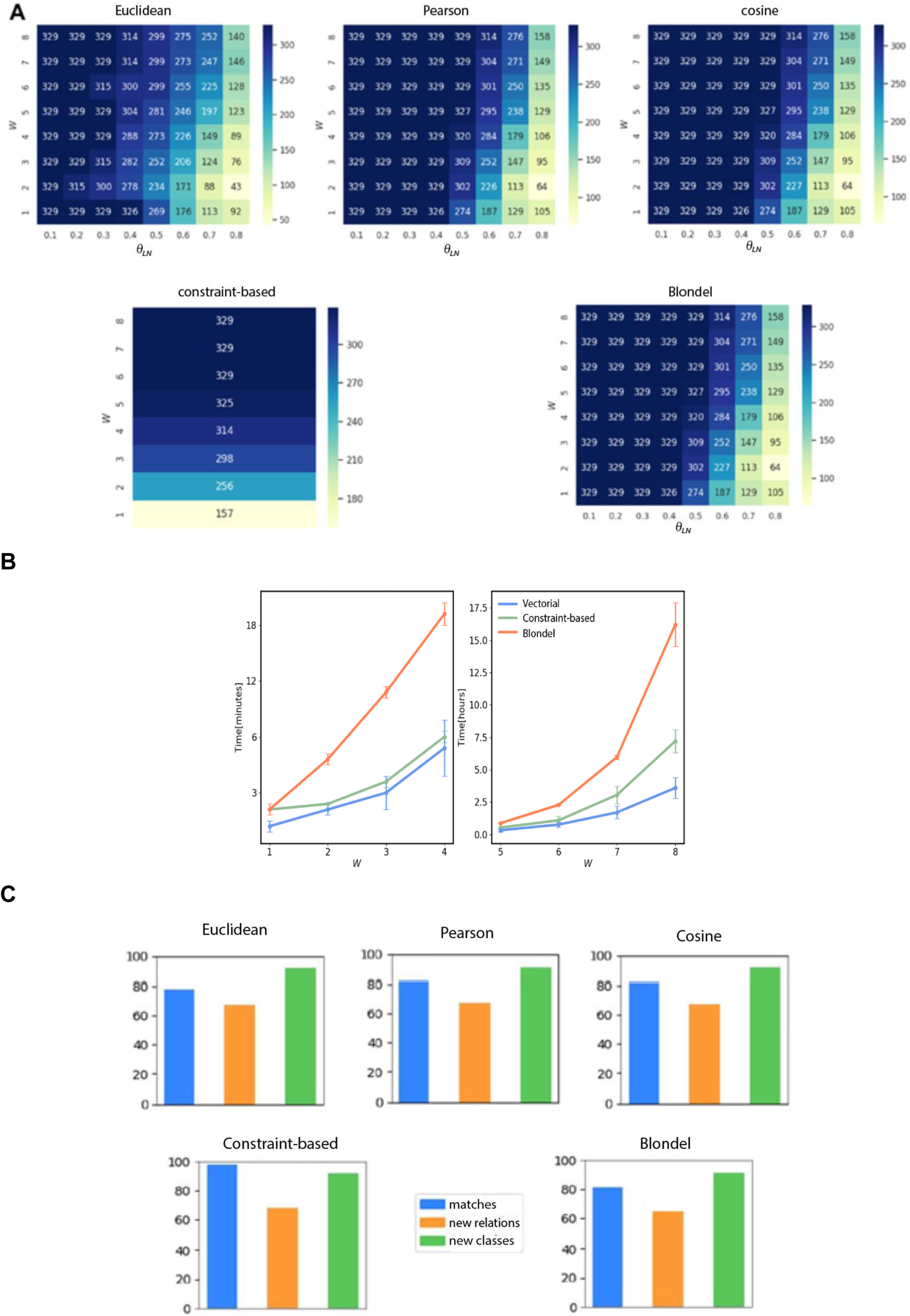
FOntCell performance merging CELDA with LifeMap. **(A)** Heat maps of the matches obtained with two-parameter combinations, window length and name score threshold, using five structure matching methods: the three vectorial structure matching Euclidean, Pearson, and cosine; constraint-based structure matching, and Blondel structure matching. The two optimized parameters are the window length *W* and the local sequence threshold *θ*_*LN*_ in the ranges [0.1, 0.8] and [1, 8], respectively, using steps of 0.1 for *θ*_*LN*_, and 1 for *W*. Bluer colour corresponds to higher number of synonyms. **(B)** Run time for the five structure-matching methods and window sizes *W* in the range [1, 8]. The vectorial structure matching {Cosine, Euclidean, Pearson} have similar runtime lines and are represented by a single line. **(C)** Percentages of matches, new classes and new relations, obtained with the five structure matching methods with merging parameters *W* = 4, *θ*_*N*_ = 0.85, and *θ*_*LN*_ = 0.7.

We observed that a name threshold *θ*_*N*_ = 0.85 assigns as similar class labels those labels that differ in orthographic variations, such as ‘s’ endings, apostrophes, etc. Therefore, we set for the remaining analysis *θ*_*LN*_ = 0.7 to be less than *θ*_*S*_ = 0.85 since we expected more name variability in nodes between subgraphs comparisons than in class-to-class comparison. We should note that we use the term node when we refer to graphs or subgraphs associated to ontologies and the term class when we refer directly to ontologies. Thus, *θ*_*LN*_ < *θ*_*N*_ recovers some meaningful cases during the structure mapping match and helps constrain the graph isomorphism problem arising during subgraph comparisons. Since smaller *W* produces smaller subgraphs, the possibility to slip into isomorph subgraphs is higher. Therefore, the topology metric is more sensitive to *θ*_*LN*_. A window *W* = 4 minimizes the sensitivity to *θ*_*LN*_ In the CELDA and LifeMap fusion, it is interesting to reduce the *θ*_*LN*_ sensitivity since a more sensitive method finds more isomorph subgraphs. If *W* is very large, the method considers very unrelated between the two graphs (corresponding to ontologies) nodes. We studied the run time of the five structure mapping methods and we found that the vectorial based methods (cosine, Euclidean and Pearson) are the faster ones and at least one order of magnitude faster that the Blondel method (Fig.1B).

To analyse the effect of each of the five structure mapping methods on the percentages of matches, new classes and new relations between them, we performed a FOntCell merging of CELDA and LifeMap for the optimized fusion parameters: *W* = 4, *θ*_*N*_ = 0.85 and *θ*_*LN*_ = 0.7, for every type of structure matching method. The results show a similar quantity of classes and relations added by the different structure matching methods, and similar number of matches (Fig.1C). The constraint-based method with *W* = 4 slips into many subgraph isomorphism, i.e., it finds too many synonyms (Fig.1A), has higher sensitivity to the change of the window size *W* than other vectorial methods. The Euclidean method is more restrictive than the other vectorial methods but more sensitive to *θ*_*LN*_ (Fig.1A). The Pearson and cosine methods are similar in the distribution of number of matches obtained for all combinations of fusion parameters (Fig.1A). The cosine method obtains exactly the same number of synonyms for all pairs of parameters (Fig.1A) and is much faster than the Blondel method (Fig.1B). Therefore, we chose to use the cosine method in the remaining analysis. For a less restrictive pair of parameters *θ*_*LN*_ = 0.1 and *W* = 1 using the cosine metric, 39.1% of classes from CELDA have a structure matching in LifeMap independently of the used structure matching method. For more restrictive parameters *θ*_*LN*_ = 0.7 and *W* = 7, we obtained 32.2% of classes with structure mapping.

The fusion of CELDA and LifeMap with *θ*_*LN*_ = 0.7 and *W* = 4 resulted in a merged ontology integrating all the 841 classes from CELDA with 567 classes from LifeMap, thus, increasing by 67.42% the cell ontology information of CELDA (Fig.2). The Interactive circular Directed Acyclic Graphs (DAGs) of CELDA, LifeMap and the resultant merged ontology are presented in Fig.3. We generate DAGs as output to visualize the input and output ontologies and illustrate which parts of the input ontologies were enriched because of the merging.

**Figure 2.**
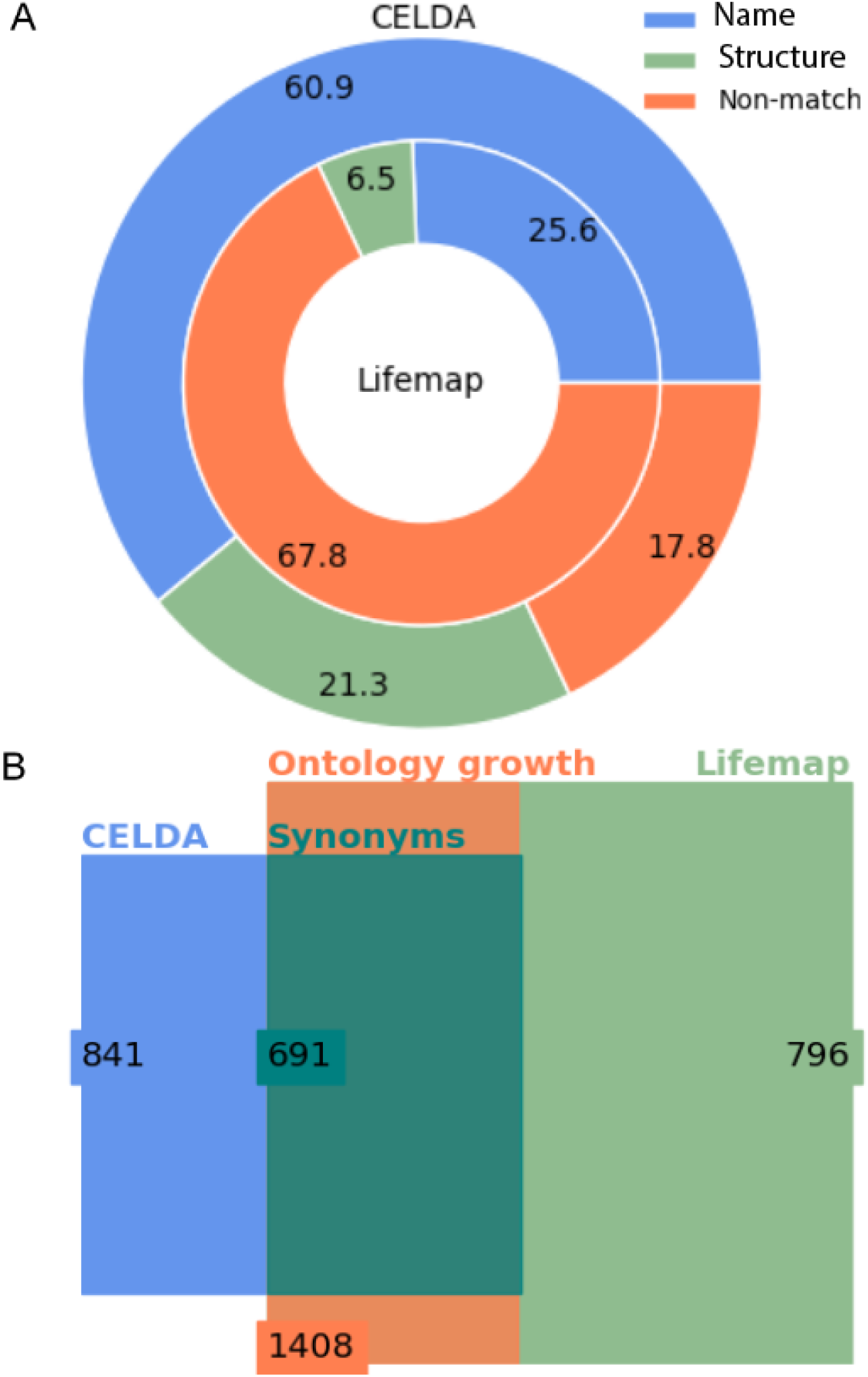
Statistics of the merging of CELDA and LifeMap with the Cosine structure matching metric and *θ*_*LN*_ = 0.7 and *W* = 4 merging parameters. (A) Donut plot of the percentages of classes added by name mapping *versus* the classes added by structure mapping to CELDA (outer circle) from LifeMap (inner circle). (B) Euler-Venn diagram with the number of classes before and after merging. The blue and light green rectangles frame the number of classes in CELDA and LifeMap, respectively, before the merging, the dark green rectangle frames the sum of name and structure equivalent classes, and the orange rectangle frames the total number of classes in the resultant CELDA and LifeMap merged ontology.

**Figure 3.**
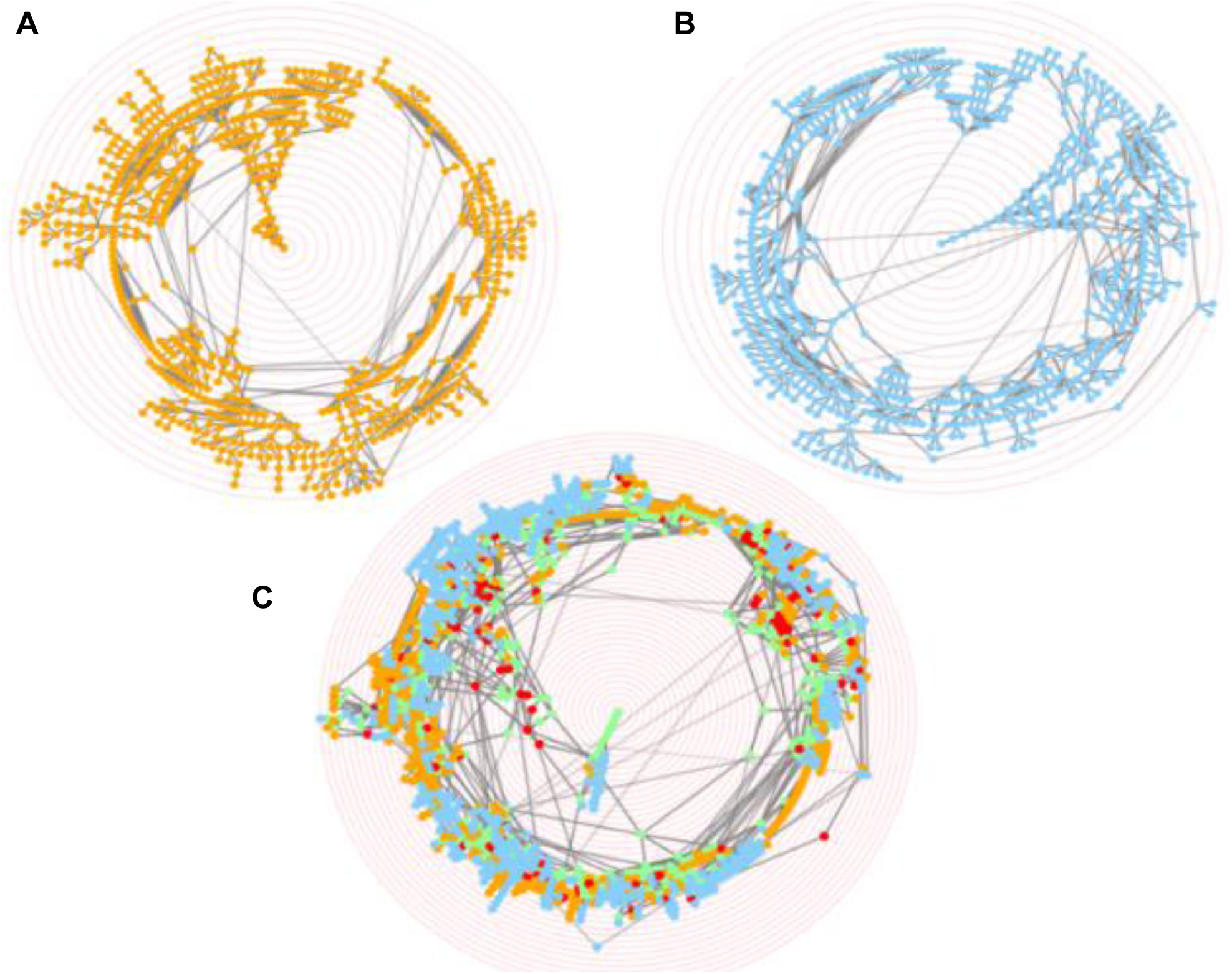
Merging of CELDA and LifeMap ontologies. Screenshots of the Interactive circular Directed Acyclic Graphs (DAGs) of (A) CELDA, (B) LifeMap and (C) the merged CELDA+LifeMap ontology, respectively. The orange and blue nodes are the non-matched contributions from ontology A and ontology B, respectively. The green and red nodes are the nodes with name and structure mapping, respectively. The ontology labels associated to the nodes appear when hovering over the nodes. The concentric red rings are zoom guides.

### FOntCell has precision of 98.63% of name mapping and mean precision of 54.84% of the five structure mapping methods when merging CELDA and LifeMap

We calculated the precision of the different mapping methods of FOntCell algorithm when merging CELDA and LifeMap with the parameters *W* = 4, *θ*_*LN*_ = 0.7 and *θ*_*N*_ = 0.85 (Fig. 4A). The results obtained through name mapping and the different structure mapping methods were validated checking manually failures (False Positive) and successes (True Positives) on the matching of the cell types. The precision was evaluated as

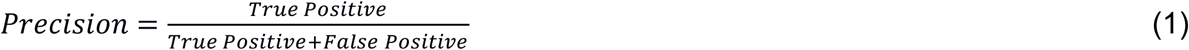

**Figure 4.**
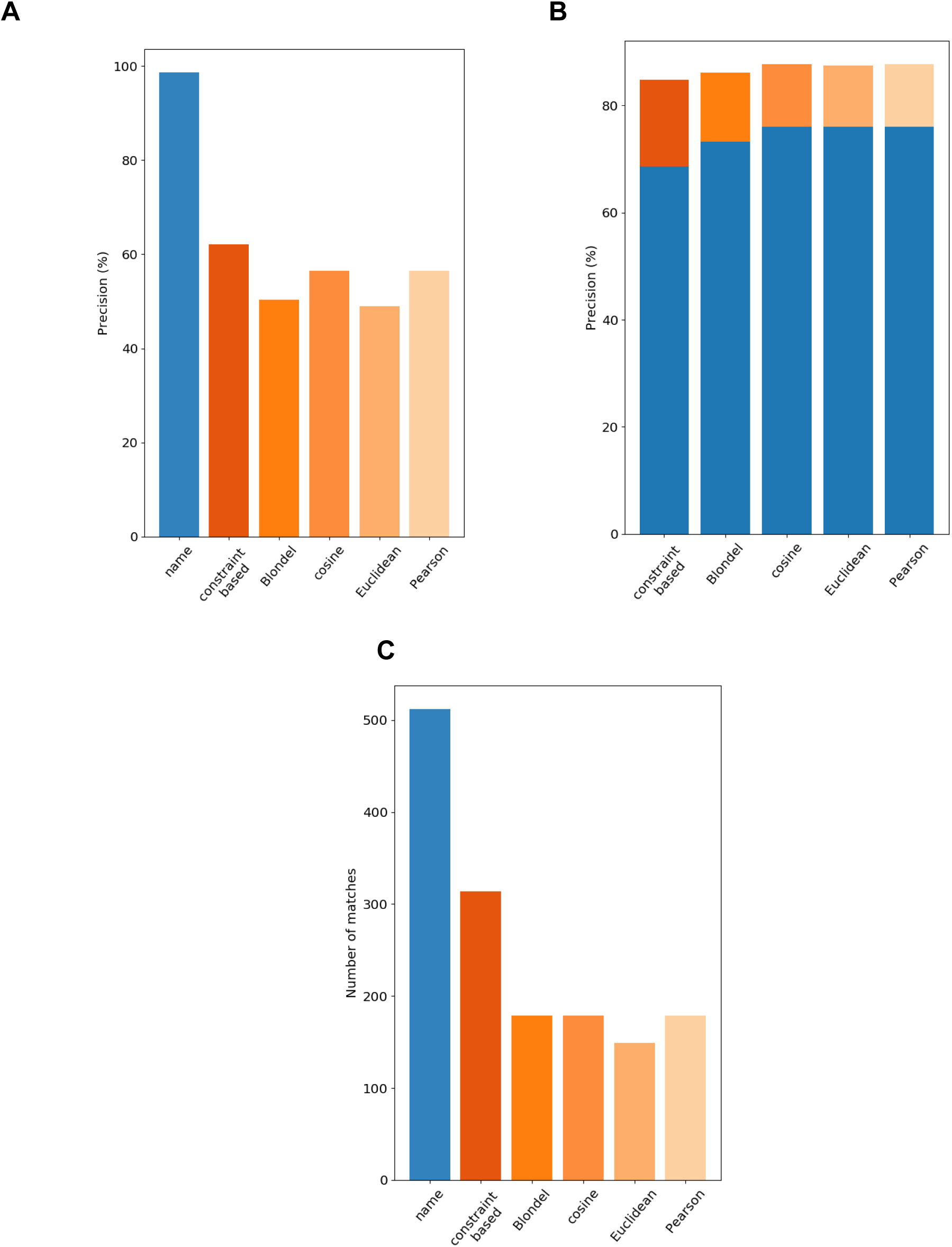
Performance of the different mapping methods of FOntCell when merging CELDA and LifeMap with parameters *W* = 4, *θ*_*LN*_ = 0.7 and *θ*_*N*_ = 0.85. (A) Precisions of name mapping and the different structure methods taken separately. (B) Combined precision of the name matching with the different structure mapping methods. The blue bars show the contribution to the precision of the name mapping, while the bars with different hues of orange show the contribution to the precision of the different structure methods. (C) Number of matches during ontology matching with the different mapping methods. Name mapping is shown in blue and the structure mappings in different hues of orange.

The name matching shows 98.63% precision (Fig. 4A) and has the highest number of matches (Fig. 4C), 512, in comparison with the other matching methods of FOntCell. Among the structure mapping methods, highest precision of 62.10% is shown by the constraint-based method, followed by the cosine and the Pearson, 56.42%, Blondel, 50.27%, and the Euclidean, 48,99%, methods (Fig. 4A). Evaluating the whole FOntCell mapping process, name and structure mapping methods taken together, we observe similar total precision with all the methods: ~ 87% using the vectorial methods, 86.1% with the Blondel method, and 86% with the constraint-based method (Fig. 4B). Anyway, total values are highest using the vectorial methods since they contribute fewer matches than the constraint-based method and the Blondel method.

When consider only structure mapping precision (Fig. 2A) the Blondel method has the second worst one, slightly higher than of the Euclidean method (Fig. 4A), however when combined with name mapping, all the vectorial methods, even then Euclidean method, surpass the precision of the Blondel method (Fig. 4B). This happens since the Blondel method has more number of matches during the structure matching than the Euclidean (Fig. 4C). This indicates that the synergies arising between name mapping and structure method combinations are stronger for the vectorial methods than for the Blondel method at least in the case of CELDA and LifeMap merging.

Considering only the structure mapping precision, the Euclidean method has the lowest one of 48.99% (Fig. 4A), while combined with name mapping it has precision of 87.44%, similar to the combined precision of the other vectorial methods, cosine and Pearson (Fig. 4B). This is due to the low number of matches obtained during structure matching, which is actually the lowest (Fig. 4C). This indicates that the synergies arising between name mapping and structure method combinations equilibrate for all the vectorial methods. The constraint-based method contributes the highest number of matches (Fig. 4C), and although it has the highest precision of 62.1% among the structure mapping methods (Fig. 4A), it decreases the total precision compared with others combined methods (Fig. 4B).

The Pearson and cosine methods show equal performance, both with the same number of matches, 179 (Fig. 4C), and the same structure mapping precision of 56.42% (Fig. 4A), which results, in combination with the name mapping, a total precision of 87.69% when combined with the name mapping (Fig. 4B). In conclusion, the cosine and Pearson methods in combination with name mapping achieve highest total precision and smallest number of matches. Therefore, we chose the cosine method as default structure matching method of FOntCell. Anyway, we could have chosen the Pearson structure method with equal results.

### The fusion of a general cell ontology CELDA+LifeMap with the LungMAP Human Anatomy (LMHA) ontology produced 65 new relations and 39 new classes related to endothelial and lymphoid cells

One of the applications of FOntCell is to merge an ontology from a broad, general description, with an ontology with very specific knowledge within the same knowledge domain. We used this functionality to merge the already merged CELDA+LifeMap with the Cell Ontology for Human Lung Maturation (LMHA) from LungMAP, i.e. a specific ontology of cells for lung development starting ~36 weeks of human fetal gestation and continuing after birth with some variation in when the alveolar stage commences and when it is complete. The .owl file used in the merging was generated by the LungMAP Consortium [U01HL122642] and downloaded from (www.lungmap.net), on April 7, 2018. The LungMAP consortium and the LungMAP Data Coordinating Center (1U01HL122638), National Heart, Lung, and Blood Institute (NHLBI). To direct the fusion, we contextualized LMHA, adding new relations to redirect the fusion. First, we deleted the classes that do not provide information about specific cell types such as the ‘immune cell’ or ‘cell type’ classes. Second, we provided some new relations and synonyms to lead the fusion. This information was written in a. txt file that allowed editing LMHA when parsed. The fusion of CELDA+LifeMap and LMHA produced 65 new relations and 39 new classes related to endothelial and lymphoid cells (Fig.5).

**Figure 5.**
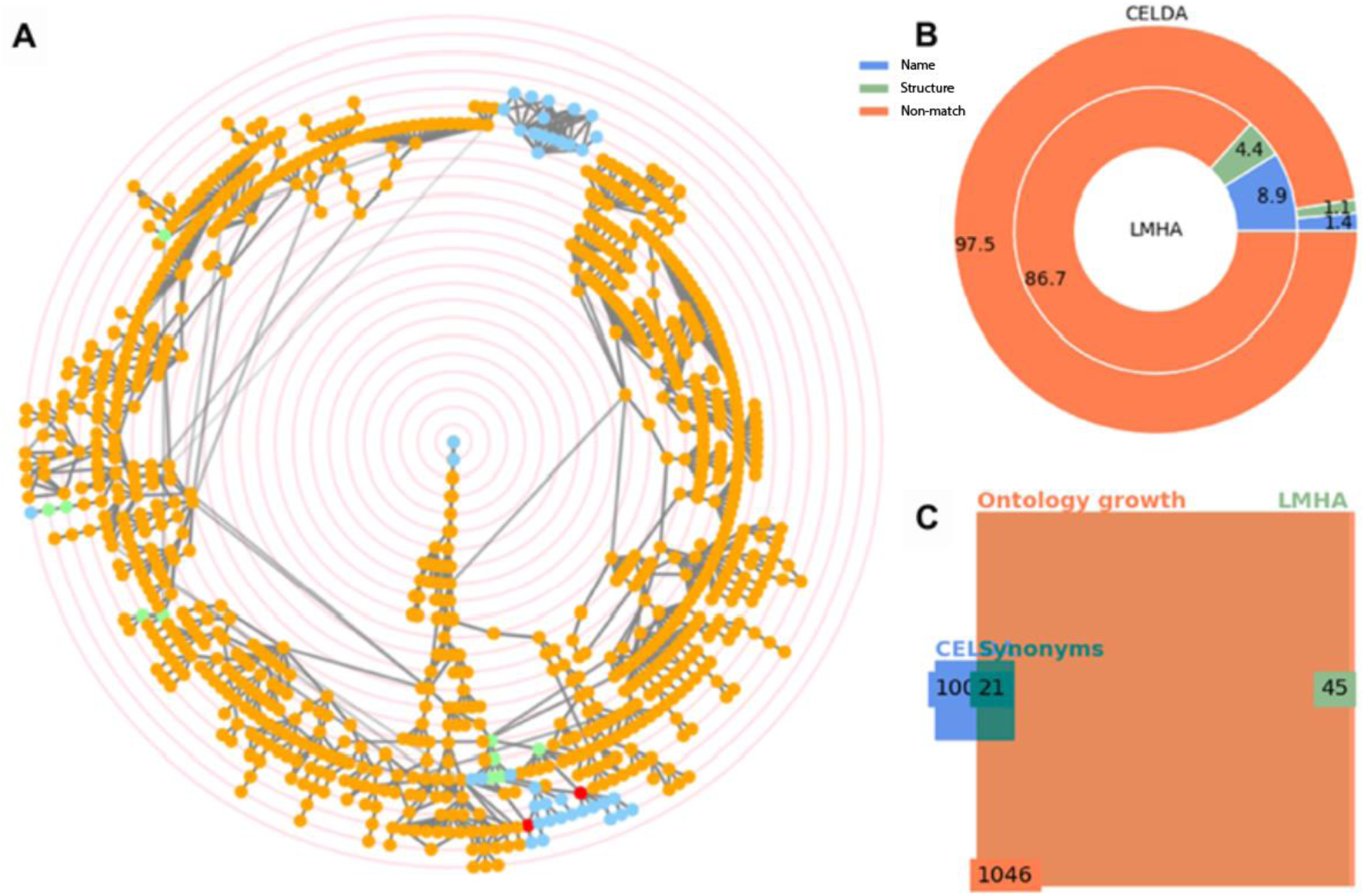
Merging of CELDA+LifeMap with LungMAP Human Anatomy (LMHA) ontology. (A) Circular Directed Acyclic Graphs (DAGs) of the merged ontology. The orange and blue nodes are the non-matched contributions from CELDA+LifeMap and LMHA, respectively. The green and red nodes are the nodes with name and structure mapping, respectively. In the interactive application generated automatically in html by FOntCell, the ontology labels associated to the nodes appear when hovering over the nodes. The concentric red rings are zoom guides. (B) Donut plot of the percentages of classes added by name mapping *versus* the classes added by structure mapping to the fused CELDA and LifeMap (outer circle) from LMHA (inner circle). (C) Euler-Venn diagram with the number of classes before and after the fusion. The blue and light green rectangles frame the number of classes in CELDA+LifeMap and LMHA before the merging, respectively, the dark green rectangle frames the sum of name and structure equivalent classes, and the orange rectangle frames the total number of classes in the resultant CELDA+LifeMap+LMHA merged ontology.

## Materials and Methods

Different ontologies use different terms to address same concepts, therefore, the first step in the fusion two or more ontologies is to align the ontologies and find the equivalent, synonymous nodes between them. Equivalent nodes are detected by a combination of text-sequence (name matching) and graph-topological similarity (structure matching) (Figure 6C). The main steps for ontology fusion are preceded by preprocessing of the ontologies: detection of equivalent nodes between the two ontologies (matching), and expansion of the non-common edges branching from the equivalent nodes (merging) (Fig. 6B). The merging works through expansion of the non-common relations branching from the matched classes (Fig. 6A). Matched classes are detected by a combination of text-sequence (name matching) and graph-topological similarity (structure matching) converting the ontologies to a graph structure according to their relations. FOntCell searches for similar (to match them) and different (to append them during the merging) classes.

**Figure 6.**
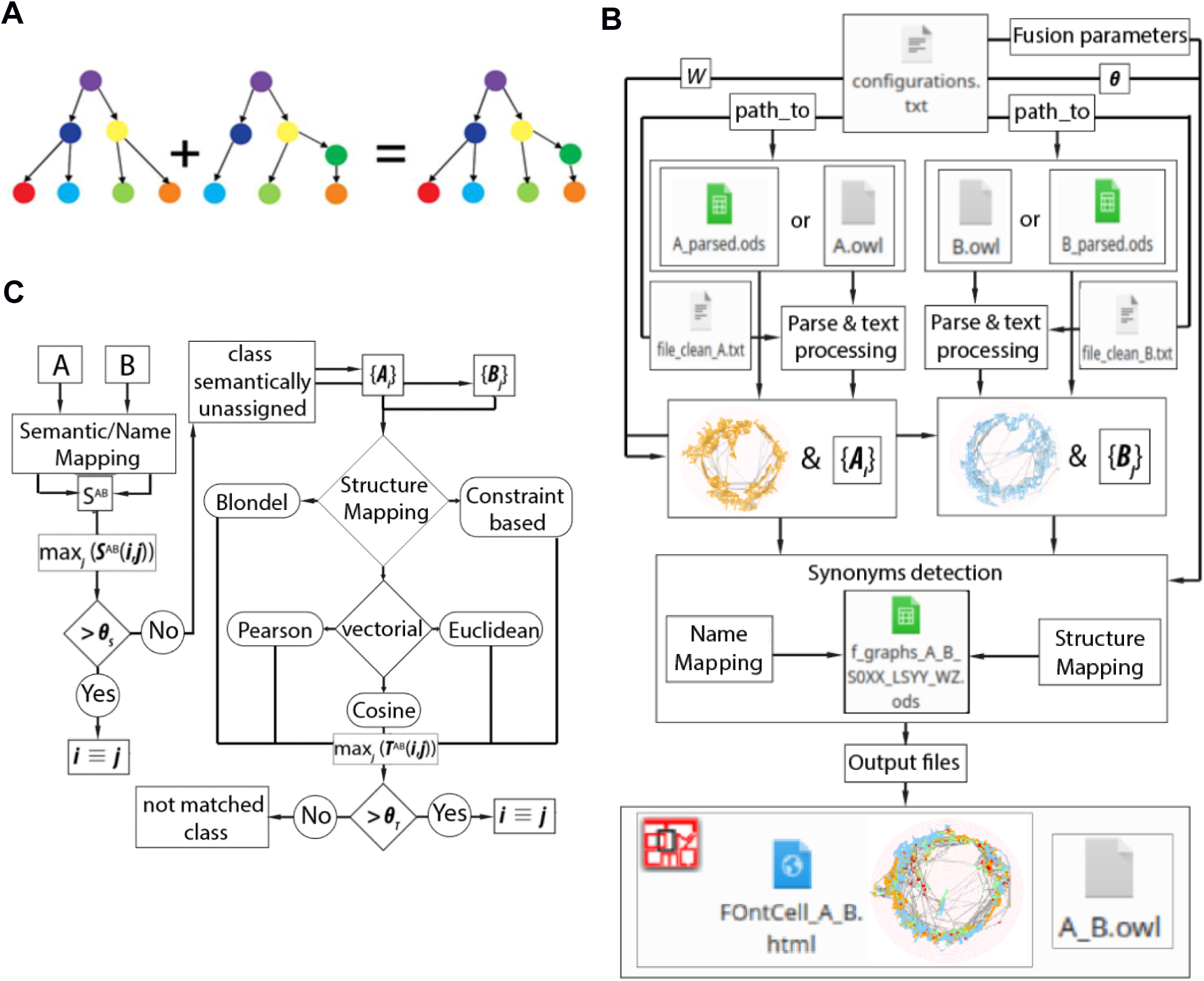
FOntCell algorithms. (A) Conceptual example of merging of ontologies. For merging two ontologies *A* and *B* into an ontology *C*, FOntCell matches first equivalent classes between *A* and *B* and then merges the non-common relations that branch from the equivalent classes. Equivalent classes are marked with same colours in the two ontologies *A* and *B*. (B) FOntCell flux diagram. **θ** is a vector with the merging parameters *θ*_*N*_, *θ*_*T*_ and *θ*_*LN*_. (C) Flux diagram of the compilation of the structure mapping for ontologies *A* and *B*. *A*_*graph*_ and *B*_*graph*_ denote the graphs of ontology A and B, respectively. {A_*i*_} and {B_*j*_} denote the sets of subgraphs around nodes *i* and *j* of ontology A and B classes, respectively. The rhombi mark two alternative decisions and the octagons three alternative processes.

### Data pre-processing and ontology parsing

FOntCell can merge any two ontologies sharing some classes and knowledge domain; however, our main interest is to apply the algorithm to the merging of cell ontologies. Different ontologies store the information in different data structures. FOntCell allows selecting as an input argument the data relation type (instances, children, parents). Since CELDA provides its cell type development information as a class attribute that has to be parsed, we implemented a pre-processing tool for parsing the cell development relations of CELDA. Additionally, the tool implements functions for class label preconditioning. In an iterative approach of use of FOntCell, the pre-processing parser reads the ontologies and selects the relations of interest (simple ascendant-descendant relations or as an argument), producing a file with parent-children relations. The parser edits automatically the ontology files to improve the fusion. Therefore, we recommend to run a ‘test’ in order to detect incorrectly fused which classes, scan useless or irrelevant information about the selected topic, and then create a. txt file with the ontology features which will be used to direct the parsing of the ontology in the next FOntCell run. Directing the parsing is a way to edit, or precondition the ontologies with FOntCell.

On one hand, CELDA uses information from other ontologies, therefore the original CELDA structure is not compact; it is split into several trees and contains information related to tissues, immortal cell lines, species, etc. Thus, FOntCell filtered CELDA by live cell-types, and by ‘human’ or ‘mouse’ species label. Additionally, in some cases CELDA does not specify the mouse-human cell origin, thus producing ‘disconnections’ in a figurative ontology associated tree. Therefore, to generate a connected cell-type development graph, we implemented a method to reconnect loose nodes (classes from ontology) to the nearest ascendant in CELDA, creating new ontology relations between classes.

On another hand, the pre-processing of LifeMap is conditioned by the fact that LifeMap is not in an OWL format, however its information about cellular type development is available at the LifeMap website repository (9). For LifeMap pre-processing we implemented a parser to substitute automatically all the symbols (’-’, ‘/’, ‘,’) in cell type labels by blank spaces. We automatically searched the LifeMap website and obtained all the information related to cell name and synonyms, development hierarchy and cell localization.

Ontologies can have redundant or missing relations. To solve such issues FOntCell can modify the original ontology relations by adding, deleting and/or fusing classes and relations with an ‘Automatic ontology editor’ that given an auxiliary. txt file that defines the disconnected/redundancy automatically edits and rewires the ontology. The format of such file is described in the FOntCell user manual. The parsing of CELDA and LifeMap generates two two-column matrices *A* and *B*, respectively, with as many rows as the number of cell type relations in the respective ontology. The first column contains the name of each class, and the second column, the name of one of its descendants. Building the matrices *A* and *B* is the first step of FOntCell in fusing the ontologies. FOntCell can merge any two ontologies in an .owl file in an OWL format, or in .ods files in parent-child relationship format compatible with the `pyexcel-ods` Python module.

### Calculation of the name-mapping matrix

FOntCell builds a name-mapping matrix measuring the similarity between the labels of two classes and, optionally, of the synonyms from the synonym attribute of two classes of different ontologies using string matching. FOntCell, among other string processing tasks, removes mismatching words, splits words, selects substrings, selects only the class name, or uses lists of synonyms representing variation of the class names.

In the simplest case of not activating the option to use synonyms, to measure the similarity between each class label of two ontologies *A* and *B*, FOntCell builds a name mapping matrix, *S*^*AB*^, using the Levenshtein metric (11), which measures the minimum number of insertions, deletions and necessary replacements to make two strings equal. To obtain the similarity in the range [0, 1], we use the opposite of the scaled Levenshtein metric:

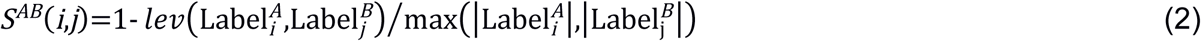

where *lev* is the Levenshtein distance between two strings. For two strings *a* and *b* of lengths |*a*| and |*b*|, respectively, the Levenshtein distance *lev*(|*a*|,|*b*|) is (11):

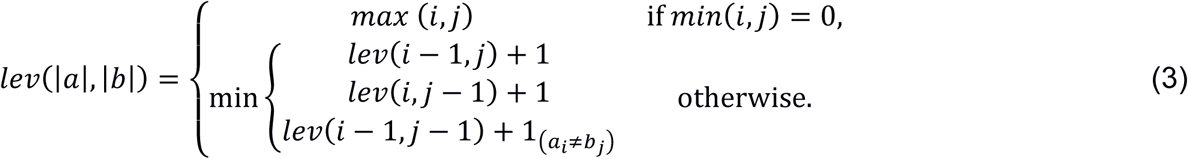

where 1_(*ai≠bj*)_ is the indicator function equal to 0 when *a*_*i*_ = *b*_*j*_, and equal to 1 otherwise, and *lev*(*i*,*j*) is the distance between the first *i* characters of *a* and the first *j* characters of *b*. Informally, the Levenshtein distance is the minimum number of single-character edits (insertions, deletions or substitutions) required to change one string into the other string. 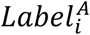 and 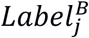 are the labels of the class *i* and *j* of the ontologies *A* and *B*, respectively, and || is the length of the string. Applying the pairwise expression (2) for each stripped label class *i* of *A*, and the stripped label class *j* of *B*, FOntCell builds the name mapping matrix *S*^*AB*^ between *A* and *B*.

In the case of selecting the option to use the synonym attribute of the classes, the similarity between two classes based on both synonym attribute and class label, {*i*}∈*A* and {*j*}∈*B*, is calculated as a submatrix of the name matching *S*^*{A}{B}*^ between each synonym of class {*i*} and each synonym of class {*j*} (including the principal label of the class) using the Levenshtein distance and taking the highest score of *S*^*{A}{B}*^ as the matching between the two classes to be used in the final name mapping matrix *S*^*AB*^(*i*,*j*) = max *S*^*{i}{j}*^. FOntCell considers that two labels have a name matching, if their score given by eq. 1 is greater than a name score threshold *θ*_*N*_ (default 0.85).

### Calculation of the structure mapping matrix

Not all classes from one ontology are identifiable as classes of the other ontology, e.g. in the CELDA and LifeMap merging, ≈ 60% of the classes from CELDA are initially not assigned to LifeMap with name mapping. FOntCell allows selecting among several types of methods to identify unmatched classes. One of the functionalities of FOntCell is to recognize matchings between two ontologies and merge them into a unique class, i.e. two labels of two classes having very different name labelling but corresponding to the same concept. FOntCell discovers synonymous classes between two ontologies using the hypothesis of structure mapping, i.e. two classes have a match if, creating a graph structure from ontology descendency, they have similar surrounding subgraphs. FOntCell measures the structure mapping between two graphs using different methods to build a structure mapping matrix *T*^*AB*^ (*a* × *b*), where *a* and *b* are the number of classes of a graph from ontology *A* and a graph from ontology *B*, and *Ã* and 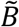 are the respective adjacency matrices, corresponding to these graph. Once a window length *W* (default 4) is selected, for each node *i* from *A* FOntCell constructs the surrounding subgraph of nodes {*i*} ∈ *A*_*w*_(*i*) and calculates its similarity with all subgraphs {*j*}∈ *B*_*w*_(*j*), where *A*_*w*_(*i*) and *B*_w_(*j*) are the subgraphs of length *W* centred in *i* and *j*, respectively. Each subgraph *A*_*w*_(*i*) is defined by a centre node *i* and all the nodes inside a window length *W* upstream or downstream of *i*. Thus, FOntCell performs a ‘structure convolutional matching’, tailoring different metrics to calculate the similarity between subgraphs *A*_*w*_(*i*) and *B*_w_(*j*).

The Blondel method initially developed to measure the similarity between graph vertices (12) can be used to assess the structure matching between two networks but is quite slow (Fig.1B). To improve the speed of the structure mapping assessment, we designed two new methods that calculate the structure matching of ontologies in a convolutional fashion: Vectorial structure matching and Constraint-based structure matching; additionally, we adapted the Blondel method to work for such new structure convolutional matching approach. An example of a convolution window sliding across a graph is depicted in Fig.7A.

**Figure 7.**
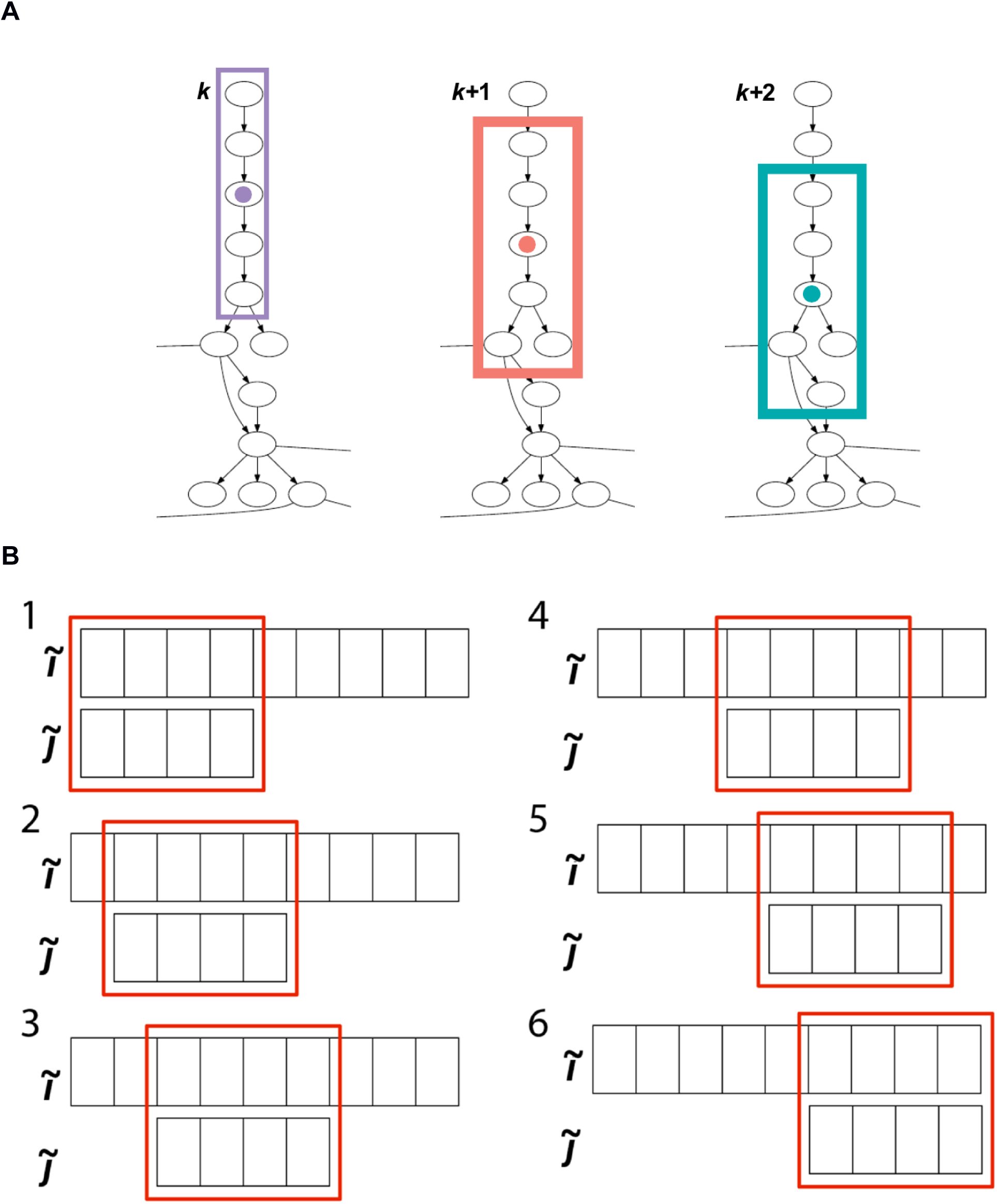
**(A)** Example of three consecutive steps of the sliding window of length *W* = 2 used in the calculation of the structure convolutional matching. For each central node, marked with a colour circle, the nodes involved in the calculation of the structure convolutional matching are framed with a rectangle of the same colour as its corresponding central node. **(B)** Example of six steps of the sliding window in the calculation of similarity between two vectors 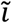 and 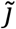 of lengths 9 and 4, respectively

### Vectorial structure matching

For each possible pair of nodes *i* ∈ *A* and *j* ∈ *B*, and window length *W*, we search for all possible nodes {*k*} ∈ *A*_*w*_(*i*) and {*l*} ∈ *B*_w_(*j*), where *A*_*w*_(*i*) and *B*_w_(*j*) are the subgraphs of length *W* centred in *i* and *j*, respectively. We take the corresponding rows 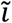 and 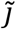 of the adjacency matrices *Ã*_*w*_(*i*) and 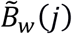. Those rows are not necessarily of the same length; therefore we calculate all the possible convolution similarities of the shorter row over the longer row (Fig.6B), using one of the metrics *M* = {1 - cosine, Euclidean, 1 - Pearson}. I.e. if *a*_*wi*_ > *b*_*wj*_, we calculate *b*_*wj*_ − *a*_*wi*_ + 1, convolution similarities *p*_*ij*_, where *a*_*wi*_ and *b*_*wj*_ are the lengths of the rows 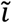 and 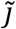 of the adjacency matrices, respectively. Then, we select the maximum similarity 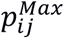. Finally, the maximum similarity 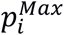 for a *j*^*Max*^ ∈ *B* across all *j* ∈ B is assigned to the structure mapping matrix *T*^*AB*^(*i*, *j*). For brevity, throughout the whole text we name the vectorial structure matching using the (1 - cosine) and (1 - Pearson) metric cosine and Pearson structure matching, respectively.

### Constraint-based structure matching

For all possible pairs of nodes *i* ∈ *A* and *j* ∈ *B*, and for a window length *W*, we search for all possible similar sequence pairs of lists of nodes {*k*} ∈ *A*_*w*_(*i*) and {*l*} ∈ *B*_w_(*j*), where *A*_*w*_(*i*) and *B*_w_(*j*) are the subgraphs of length *W* centred in *i* and *j*, respectively. Then, for each node *k* in the list {*k*} we calculate the shortest path *s*_*ki*_ to *i*, and we produce a list {*s*_*ki*_} of shortest paths. Next, we assign to each *s*_*ki*_ a constraint value *c*_*ki*_ = *W* + 1 − *s*_*ki*_. Finally, we sum the list {*c*_*ki*_} to produce an accumulated constraint *C*_*i*_ and assign it to the structure mapping matrix *T*^*AB*^(*i*, *j*).

### Blondel structure matching

FOntCell uses the Blondel metric

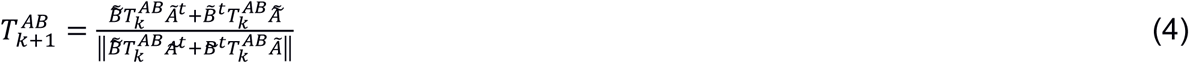

where *t* is the transpose operator and eq. 4 is calculated iteratively until an even number of steps *k* of convergence to a stable structure matching *T*^*AB*^. For each node *i* from *A*, FOntCell constructs the surrounding subgraph {*i*}∈*A* and calculates its similarity with all subgraphs {*j*}∈ *B* of *B* using eq. 4 with the adjacency matrices of each subgraph. FOntCell performs a structure convolution, tailoring eq. 4 to the case of subgraphs {*i*} and {*j*}

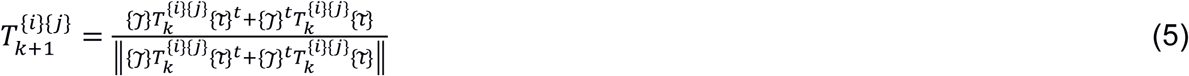

where 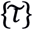 and 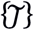 are the adjacency matrices of the respective subgraphs {*i*} and {*j*}. Finally, the structure score on the position of *i* and *j* in the 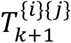 matrix is assigned to *T*^*AB*^(*i, j*).

Each of the above defined structure matching methods returns a structure score between two nodes (*i*, *j*) defined by *T*^*AB*^(*i*, *j*) where *i* ∈ *A* and *j* ∈ *B*. This convolution improves the result of the structure mapping over the whole graphs since it reduces the influence of distant nodes and edges. Name mapping and structure mapping carry complementary information and FOntCell regains information from both.

### Ontology matching

To match classes, FOntCell initially selects the best match for each node *i* from *A* with a node *j* from *B*, using the name mapping matrix *S*^*AB*^. If *S*^*AB*^(*i*,*j*) > *θ*_*N*_, FOntCell considers classes *i* and *j* as matched and classifies this assignment as a ‘name match’. If *S*^*AB*^(*i*,*j*) ≤ *θ*_*N*_, FOntCell takes the element *T*^*AB*^(*i*, *j*) from one of the aforementioned structure matching methods selected by the user to calculate the structure mapping and considers the nodes *i* and *j* matched if *T*^*AB*^(*i*, *j*) ≥ *θ*_*T*_, where *θ*_*T*_ is a structure mapping threshold selected by the user. To improve the result achieved with the structure matching method, FOntCell performs a further local name comparison using the name mapping matrix to calculate the mean of the name match *S*^{*i*}{*j*}^ of each subgraph pair {*i*} {*j*} built around the central nodes *i* and *j*, and the same window size *W* used to calculate *T*^{*i*}{*j*}^. FOntCell takes the best name scores from *S*^*AB*^(*i*, *j*), calculates the mean of these name matching scores for the pair {*i*} {*j*}, and then builds a new name matching matrix of {*i*} {*j*}: *S*^{*i*}{*j*}^. If *S*^{*i*}{*j*}^ > *θ*_*LN*_, where *θ*_*LN*_ is a local name matching threshold (default value *θ*_*LN*_ = 0.7), FOntCell considers nodes *i* and *j* as synonyms and classifies the corresponding classes as a ‘structure match’ (see Fig.5C). FOntCell creates a file with the relevant information about each node from *A*: with columns (1) native node label in *A*, (2) translated node label assigned from *B*, (3) name score, (4) structure score and (5) type of assignment’ (Name/Structure). In case of no assignment, the ‘type of assignment’ is marked as ‘Non-matched’.

### Ontology merging

Once the matched classes between two ontologies are detected, FOntCell translates the name-labels of all classes from ontology *B* to their equivalent names, if any, in ontology *A*. Next, FOntCell appends the translated classes from *B* to *A*. Then, it performs an ordered-set operation to eliminate all the possible class-relations repeats generated from the appendage. The resulting relation array represents the merging of the two ontologies. In addition, FOntCell creates an OWL format file with the result of the fusion by reading the .owl file of ontology A and appending the new classes from B at the start of the ontology class site. The information stored in these new classes is: (1) new ID, (2) class label, (3) class synonyms, and (4) ascendant relationship. Finally, FOntCell creates an. html file with an interactive circular Directed Acyclic Graph (DAG) of the original and fused ontologies, and statistical information of the fusion, i.e. percentage and number of added classes/relations and type of matches in textual and graphical form.

## Implementation

FOntCell is developed in Python v3.7 and uses the Python library `NetworkX` to derive the digraph relation of the ontology and to transform each class to a node and each hierarchy step to an edge. `NetworkX` graphs allows FOntCell access the sorted list of nodes without repeats, and produce digraphs compatible with graph visualization tools such as `graphviz` and `matplotlib`. For specific data manipulation, FOntCell uses `numpy`, `pyexcell_ods`, `argparse`, `stringdist`, and basic Python libraries such as `os`, `collections` and `itertools`. The algorithm complexity (Big O) is quadratic time O(n^2^). For parallelization and the structure mapping, FOntCell uses BigMPI4py (13). We added a demo function to the FOntCell distribution package merging CELDA with LifeMap.

The automatic installation installs all the dependencies. If some issues occur during installation or run, please see the title page at https://pypi.org/project/fontcell/. Full instructions of the prerequisites for installation, the downloading of FOntCell, the user manual, an example of use of how to run FOntCell and an example of the html output created by FOntCell are provided in the Supplementary material file.

## Software Availability

The FOntCell software package, written as a Python module is available at https://pypi.org/project/fontcell/, and the resulting fusion of CELDA and LifeMap cell ontologies in OWL format at https://gitlab.com/JavierCabau/fontcell

## Funding

J C-L has been supported by a FPI grant of the Ministry of Economy and Competitiveness, Spain, BES-2017-080625. DG and MJ A-B. have been supported by grants DFG113/18 from Diputaci6n Foral de Gipuzkoa, Spain, Ministry of Economy and Competitiveness, Spain, MINECO grant BFU2016-77987-P and Instituto de Salud Carlos III (AC17/00012) co-funded by the European Union (Eracosysmed/H2020 Grant Agreement No. 643271).

## Conflict of Interest

none declared.

## Supplementary material

**FOntCell** is a software module in Python for automatic computed fusion of ontologies. **FOntCell** produces the results of the merged ontology in 0B0 format that can be iteratively reused by F0ntCell to adapt continuously the ontologies with the new data, such of the Human Cell Atlas, endlessly produced by data-driven classification methods. To navigate easily across the fused ontologies, it generates HTML files with tabulated and graphic summaries, and an interactive circular Directed Acyclic Graphs of the merged results.

This document contains:

- The prerequisites for installation of **FOntCell**.
- The instructions to download **FOntCell**.
- The User manual of **FOntCell**.
- Example of how to run **FOntCell**.
- Example of the html output created by **FOntCell**.

### Prerequisites for installation of FOntCell

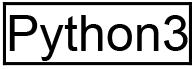

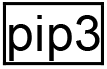

It is necessary to use pip3 to download **FOntCell** and use it with python3.

To install pip3 over Python3 use the command:

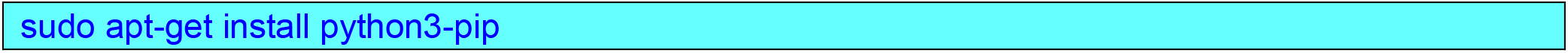

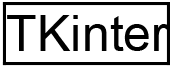

Tkinter library is a dependency that will not be installed during **FOntCell** installation.

If TKinter has not been previously installed, an error will occur during **FOntCell** import.

To install TKinter over Python3 use the command:

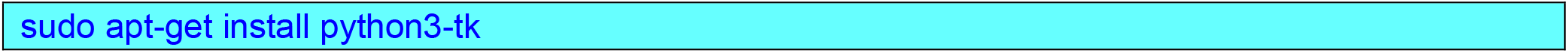

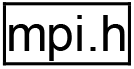

Mpi.h is an mpi4py dependency that might not have been installed during mpi4py installation, and if it is had not been installed, an error might occur.

To install mpi.h use the command:

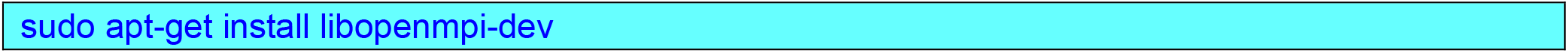

### Instructions to download FOntCell

**FOntCell** module is available at PyPI and can be installed using the command:

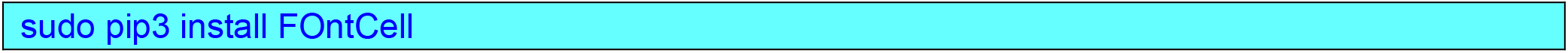

### User manual of FOntCell

After downloading and installing **FOntCell** using pip3, the user has to create the following directories:

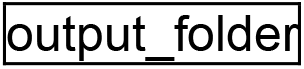

Is the directory where **FOntCell** will output the results:

- Fused ontology file
- html files
- Figures

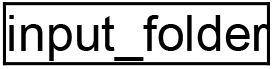

The input_folder should contain a Configuration file, the two ontologies, and the ontology_edit documents, that should be placed there by the user.

#### Files required in the Input directory

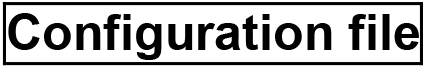

The configuration file should have the following arguments in. txt format.

*Most arguments will be for ontology 1 (A) and ontology 2 (B)*

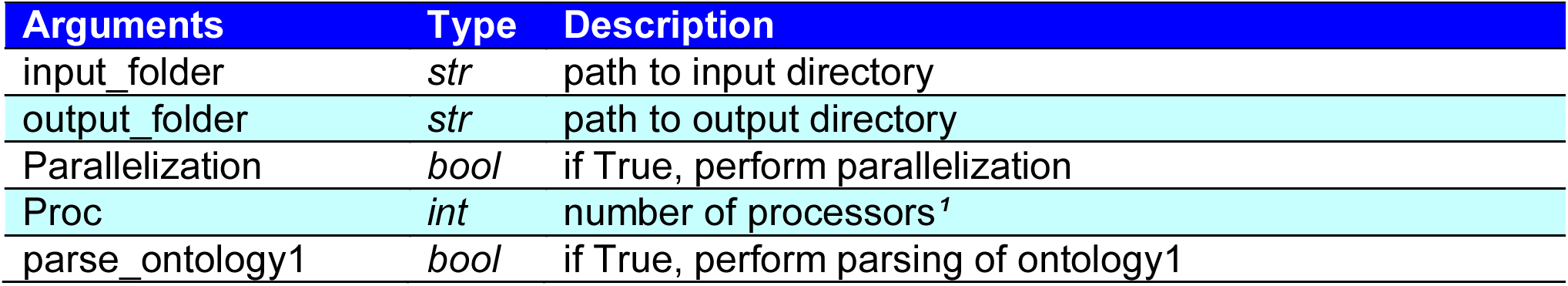
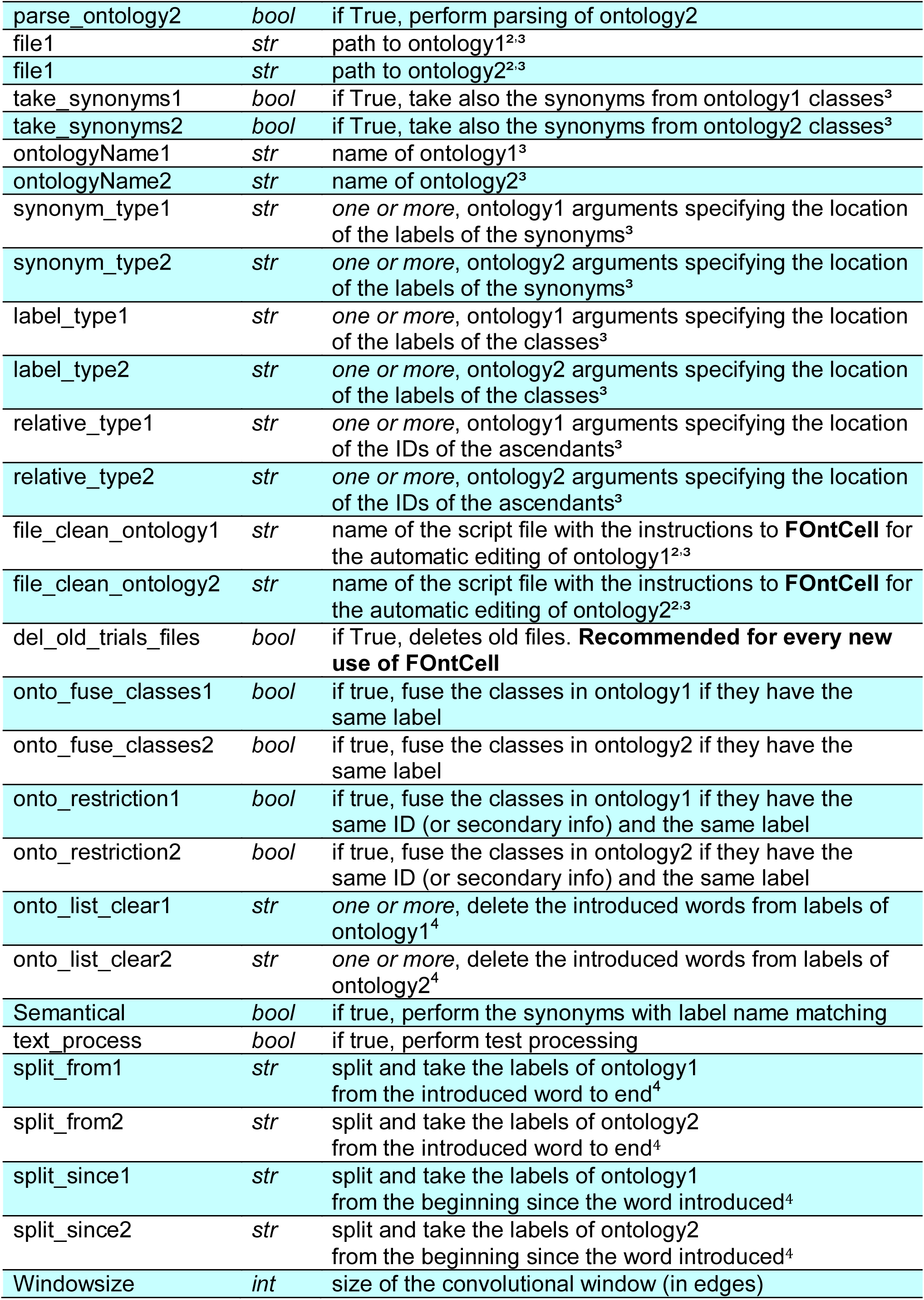
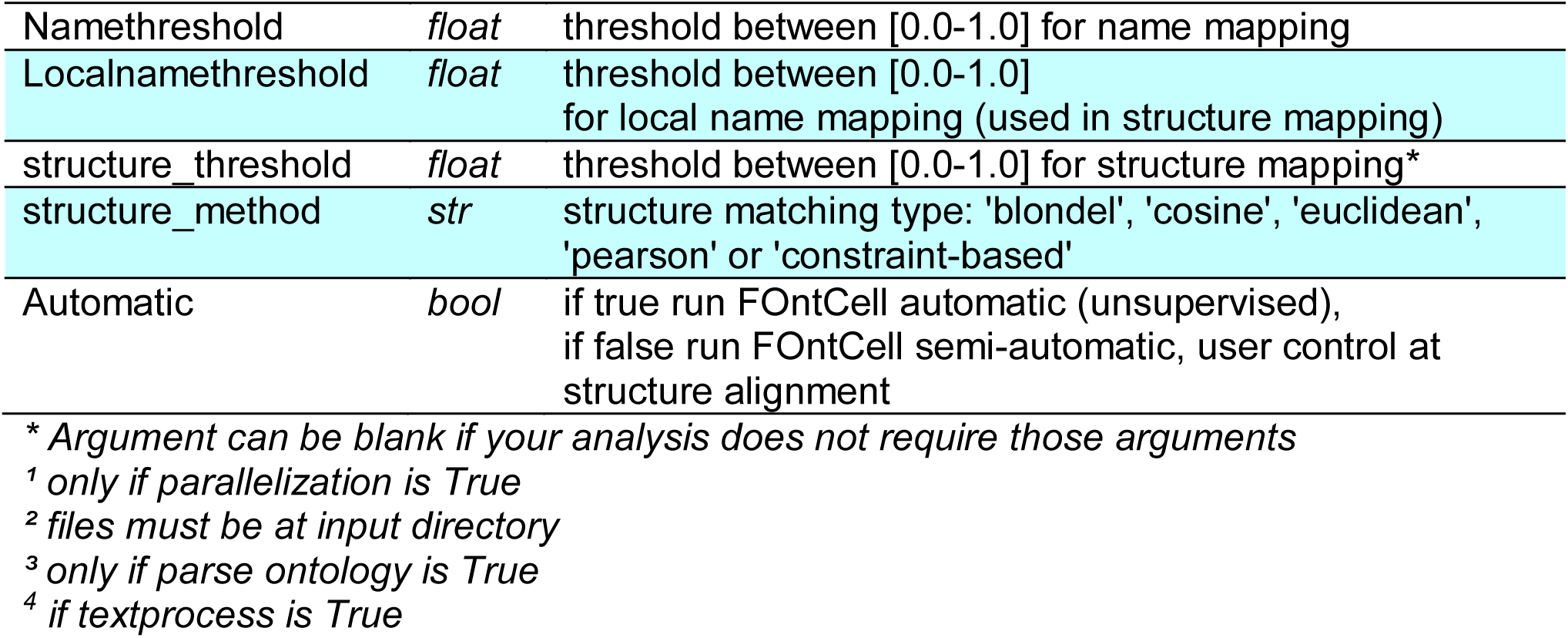

| Arguments | Type | Description |
| --- | --- | --- |
| `input_folder` | *str* | path to input directory |
| `output_folder` | *str* | path to output directory |
| `Parallelization` | *bool* | if True, perform parallelization |
| `Proc` | *int* | number of processors<sup>1</sup> |
| `parse_ontology1` | *bool* | if True, perform parsing of ontology1 |
| parse_ontology2 | *bool* | if True, perform parsing of ontology2 |
| file1 | *str* | path to ontology1<sup>2,3</sup> |
| file2 | *str* | path to ontology2<sup>2,3</sup> |
| take_synonyms1 | *bool* | if True, take also the synonyms from ontology1 classes<sup>3</sup> |
| take_synonyms2 | *bool* | if True, take also the synonyms from ontology2 classes<sup>3</sup> |
| ontologyName1 | *str* | name of ontology1<sup>3</sup> |
| ontologyName2 | *str* | name of ontology2<sup>3</sup> |
| synonym_type1 | *str* | *one or more*, ontology1 arguments specifying the location of the labels of the synonyms<sup>3</sup> |
| synonym_type2 | *str* | *one or more*, ontology2 arguments specifying the location of the labels of the synonyms<sup>3</sup> |
| label_type1 | *str* | *one or more*, ontology1 arguments specifying the location of the labels of the classes<sup>3</sup> |
| label_type2 | *str* | *one or more*, ontology2 arguments specifying the location of the labels of the classes<sup>3</sup> |
| relative_type1 | *str* | *one or more*, ontology1 arguments specifying the location of the IDs of the ascendants<sup>3</sup> |
| relative_type2 | *str* | *one or more*, ontology2 arguments specifying the location of the IDs of the ascendants<sup>3</sup> |
| file_clean_ontology1 | *str* | name of the script file with the instructions to **FontCell** for the automatic editing of ontology1<sup>2,3</sup> |
| file_clean_ontology2 | *str* | name of the script file with the instructions to **FontCell** for the automatic editing of ontology2<sup>2,3</sup> |
| del_old_trials_files | *bool* | if True, deletes old files. **Recommended for every new use of FontCell** |
| onto_fuse_classes1 | *bool* | if true, fuse the classes in ontology1 if they have the same label |
| onto_fuse_classes2 | *bool* | if true, fuse the classes in ontology2 if they have the same label |
| onto_restriction1 | *bool* | if true, fuse the classes in ontology1 if they have the same ID (or secondary info) and the same label |
| onto_restriction2 | *bool* | if true, fuse the classes in ontology2 if they have the same ID (or secondary info) and the same label |
| onto_list_clear1 | *str* | *one or more*, delete the introduced words from labels of ontology1<sup>4</sup> |
| onto_list_clear2 | *str* | *one or more*, delete the introduced words from labels of ontology2<sup>4</sup> |
| Semantical | *bool* | if true, perform the synonyms with label name matching |
| text_process | *bool* | if true, perform test processing |
| split_from1 | *str* | split and take the labels of ontology1 from the introduced word to end<sup>4</sup> |
| split_from2 | *str* | split and take the labels of ontology2 from the introduced word to end<sup>4</sup> |
| split_since1 | *str* | split and take the labels of ontology1 from the beginning since the word introduced<sup>4</sup> |
| split_since2 | *str* | split and take the labels of ontology2 from the beginning since the word introduced<sup>4</sup> |
| Windowsize | *int* | size of the convolutional window (in edges) |
| Namethreshold | *float* | threshold between [0.0-1.0] for name mapping |
| Localnamethreshold | *float* | threshold between [0.0-1.0] for local name mapping (used in structure mapping) |
| structure_threshold | *float* | threshold between [0.0-1.0] for structure mapping* |
| structure_method | *str* | structure matching type: 'blondel', 'cosine', 'euclidean', 'pearson' or 'constraint-based' |
| Automatic | *bool* | if true run FOntCell automatic (unsupervised), if false run FOntCell semi-automatic, user control at structure alignment |

\* *Argument can be blank if your analysis does not require those arguments*
<sup>1</sup> *only if parallelization is True*
<sup>2</sup> *files must be at input directory*
<sup>3</sup> *only if parse ontology is True*
<sup>4</sup> *if textprocess is True*

Every argument of the configuration.txt file needs to be precede by a a ‘>’ character forward to be parsed.

The user does not have to introduce a threshold for a test that it has not been selected

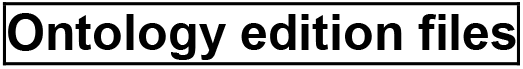

These files allows the user to edit the ontology after parse in order to direct the ontology fusion.

The document must be in. txt file with the following instructions defining how to automatic edit the ontology file:

| Instruction | Syntax | Description | Example |
| --- | --- | --- | --- |
| delete class | '...' | deletes the nodes/classes that contains the label | 'ontology label x' |
| concatenate classes | [[...], [...]] | concatenates two classes, allows introduce new classes | [[[ontology_class_synonyms]1, ID:0],<br>[[ontology_class_synonyms2], ID:1]] |
| delete class from ID | ID: '...' | deletes the class that has the ID introduced | ID:'xxxxx' |
| fuse classes | f[[...], [...]] | fuses two classes. The resultant class conserves the ID from the first class, and the descendant-ascendant relations) | f[[ontology_synonyms1],<br>[ontology_synonyms2]] |

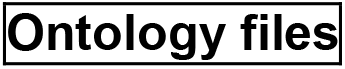

The ontologies to be fused can be in .owl format (that requires a parse) or in a .ods format, if one wishes to save the parse step.

The .ods file has the graph-edge information in two columns as in the following example:

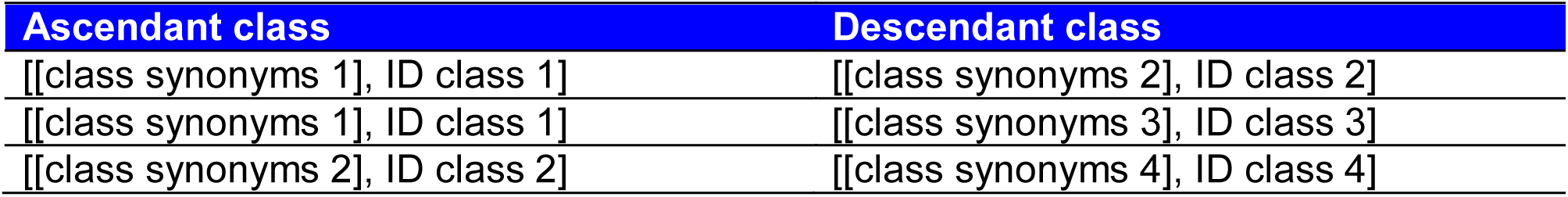

The first column corresponds to ‘parent’ and the second column to the ‘descendant’. The possible synonyms of each class are separated by commas.

#### Files created in the Output directory after fusion

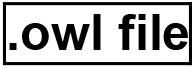

This document is the resultant ontology from the fusion of two ontologies. The structure is the same as of ontology 1 (A), with the new classes added from ontology 2 (B) at the top of ontology class section.

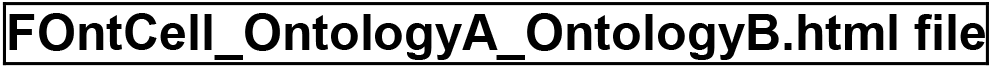

This file contains information about the fusion. First, one can see three interactive circular graphs: ontology A, ontology B and the merged ontology. Additionally, it contains other information such as: the different thresholds values and statistics about the fusion. The file also shows a direct link to the 0B0-format merged ontology, and a representative image of type of node assignation between ontology A and B (a donut graph) and an image of an Euler-Venn diagram (using squares) about how the fusion has worked.

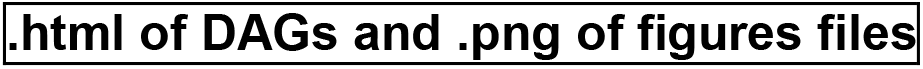

These files are the files encrusted in the F0ntCell_0ntologyA_0ntologyB.html file. The. html files are all the different interactive circular graphs, and the. png files contain the Euler-Venn diagram and the donut graph.

### Troubleshooting

For parallel computation **FOntCell** requires bigmpi4py: (https://www.biorxiv.org/content/10.1101/517441v1)

If bigmpi4py has been installed in a different conda environment from the one of the **FOntCell** installation, the parallelization will not work well (all the processes will run on a single processor). In this case, **FOntCell** will work but without parallelization.

Running **FOntCell** will raise a problem if graphviz has not been properly installed. For a correct graphviz installation, type in the command line:

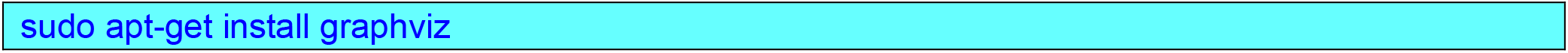

### Example of how to run FOntCell

Open python3 in bash:

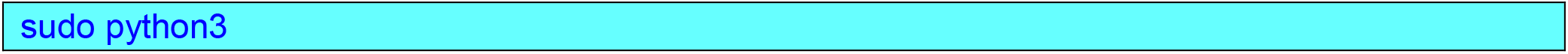

Import the **FOntCell** module:

import FOntCell

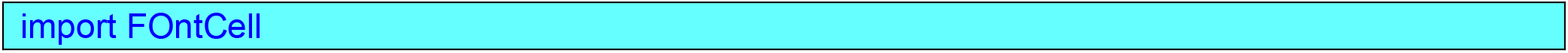

After the **FOntCell** module is imported, one can use the following functions:

To run the fusion of two ontologies:

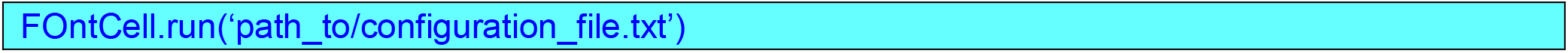

To run the demo of the fusion of CELDA with Lifemap:

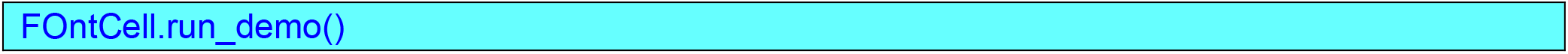

To clean internal files from old runs (recommended for use before a new fusion, especially if one of the ontologies from a previous fusion will be used again):

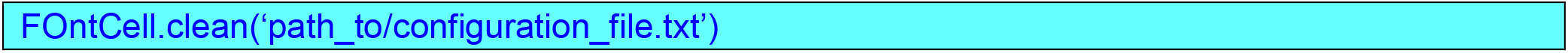

### Example of the html output created by FOntCell

In the next pages is attached the html file resulting of using **FOntCell** to merge CELDA and LifeMap ontologies to produce comprehensive cell ontology.

## FOntCell Fusion of CELDA and LifeMap

### Interactive circular Directed Acyclic Graphs (DAGs) of (a) CELDA, (b) LifeMap and (c) merged ontologies

<u>(a) DAG of CELDA ontology classes (nodes in orange)</u>

<u>(b) DAG of LifeMap ontology classes (nodes in blue)</u>

<u>(c) DAG of the Fused ontology classes (nodes in orange (from CELDA), blue(from LifeMap), red for structure match and green for name match)</u>

The ontology labels associated to the classes appear when hovering over the nodes.

Some nodes may appear overlapping.

### Parameters of the FOntCell fusion algorithm

- Name matching threshold Θ_S_: 0.85
- structure matching method: cosine
- Local Name matching threshold Θ_SL_: 0.7
- Structure matching threshold Θ_T_: 0.0

### Statistics of the input ontologies

- Number of classes of CELDA ontology: 841
- Number of relations between classes of CELDA ontology: 966
- Number of classes of LifeMap ontology: 796
- Number of relations between classes LifeMap ontology: 924

### Statistics of the merged ontology

#### Statistics of the merged by name mapping

- Number of classes with equivalence found in CELDA by name mapping: 512
- Number of classes with equivalence found in LifeMap by name mapping: 204
- Percentage of classes (in relation to the number of classes of CELDA ontology) added to CELDA by name mapping: 60.88%
- Percentage of nodes added to LifeMap (in relation to the number of nodes of LifeMap ontology) by name mapping: 25.63%

#### Statistics of the fusion by structure mapping

- Number of classes with equivalence found in CELDA by structure mapping: 179
- Number of classes with equivalence found in LifeMap by structure mapping: 52
- Percentage of classes added to CELDA (in relation to the number of classes of CELDA ontology) by structure mapping: 21.28%
- Percentage of classes added to LifeMap (in relation to the number of classes of LifeMap ontology) by structure mapping: 6.53%

#### Statistics of the fusion of non-matched nodes

- Number of classes in CELDA non-matched in LifeMap: 150
- Percentage of classes in CELDA non-matched in LifeMap (in relation to the number of classes of CELDA ontology): 17.84%
- Number of classes in LifeMap non-matched in CELDA: 540
- Percentage of classes in LifeMap non-matched in CELDA (in relation to the number of classes of LifeMap ontology): 67.84%

#### Statistics of the fusion by name and structure mapping

- Number of classes added in total (by name mapping and by structure mapping): 567
- Percentage of classes added in total (by name mapping and by structure mapping): 67.42%
- Number of relations between classes added in total (by name mapping and by structure mapping): 890
- Percentage of relations between classes added in total (by name mapping and by structure mapping): 92.13%

Added classes refers to the descendants classes founded on the mapping

### Merged ontology in OBO format

Merged ontology from CELDA and LifeMap:

<u>here</u>

### Results of the merged ontology

#### Percentages of contribution of classes to the merged ontology in relation to the classes of each contributing ontology

##### Outter circle: Numbers and percentages of CELDA

- Blue: Classes with name match: 512, percentage: 60.88
- Green: Classes with structure match: 179, percentage: 21.28
- Orange: Non-matched classes: 150, percentage: 17.84

##### Inner circle: Numbers and percentages of LifeMap

- Blue: Classes with name match: 204, percentage: 25.63
- Green: Classes with structure match: 52, percentage: 6.53
- Orange: Non-matched classes: 540, percentage: 67.84

##### Euler-Venn diagram of the classes of CELDA and LifeMap merging

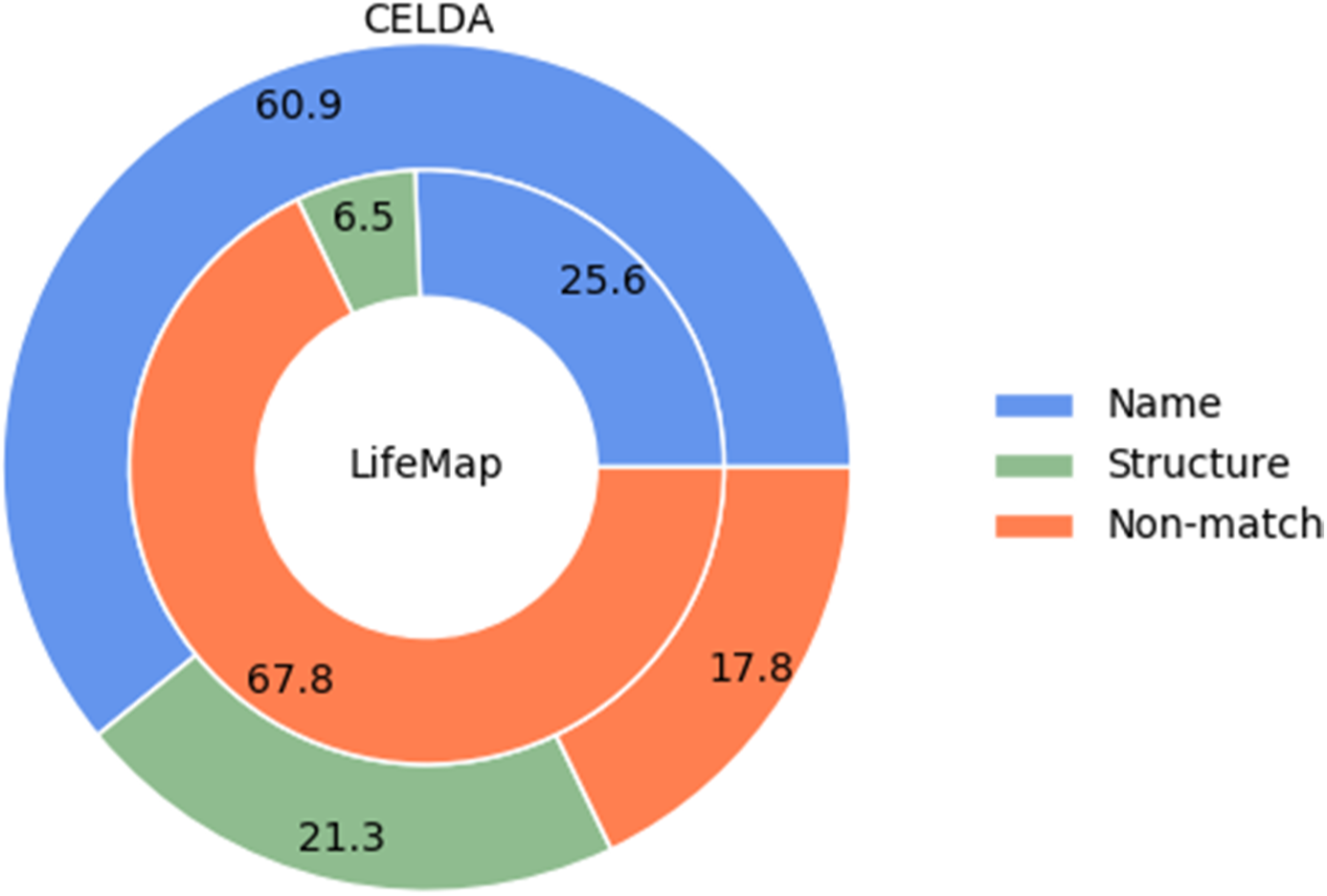

- Classes from CELDA: 841 (blue)
- Classes from LifeMap: 796 (green)
- Synonyms found in CELDA: 691 (Blue-green)
- Resulted ontology classes: CELDA classes: 841 + added classes: 567

### Additional results

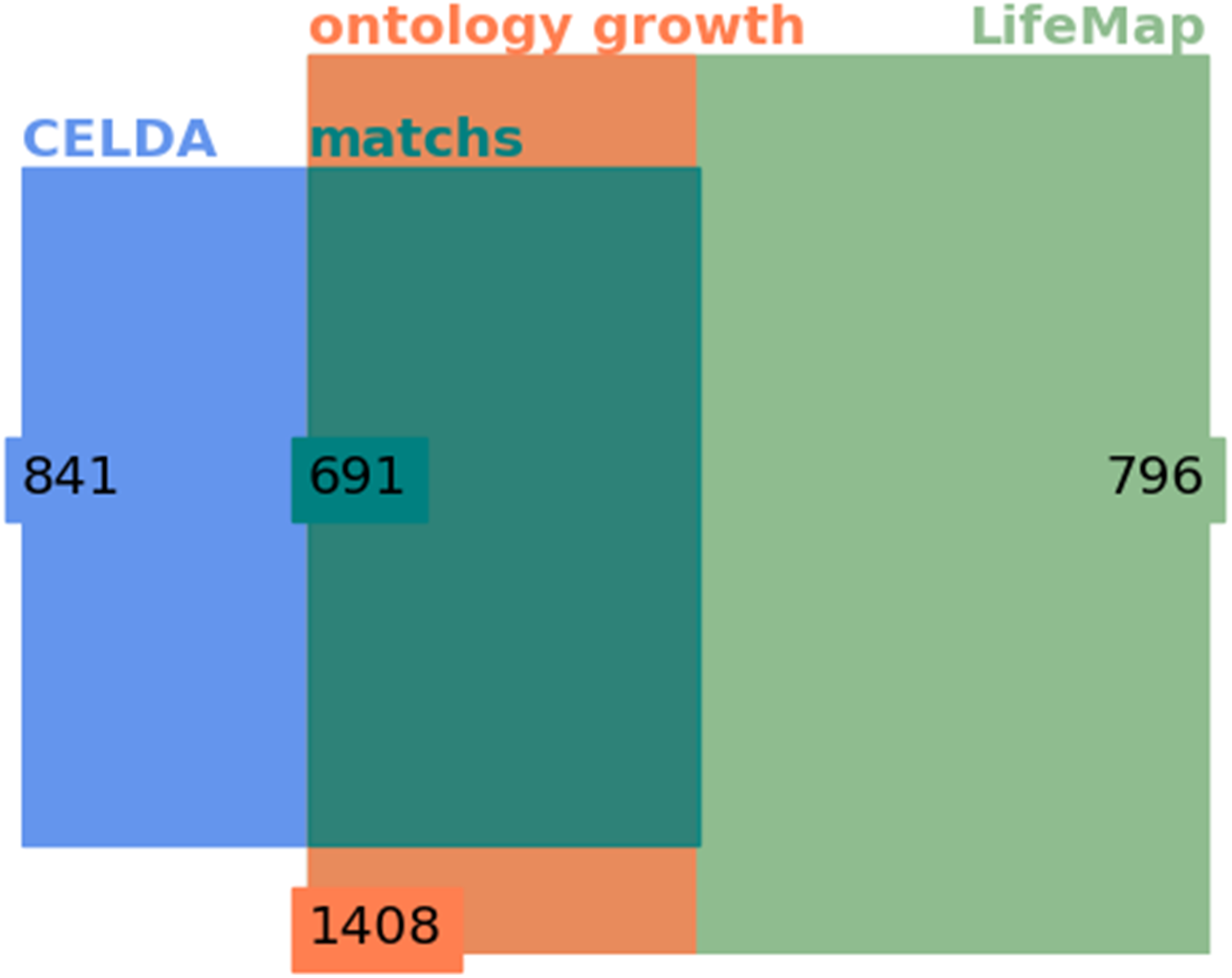

Files with results on detection of matchs, merging and name matching matrix are available at: /usr/local/lib/python3.6/dist-packages/F0ntCell/fontcell_files/

